# The effect of UVA light/8-methoxypsoralen exposure used in Extracorporeal Photopheresis treatment on platelets and extracellular vesicles

**DOI:** 10.1101/2023.10.18.562996

**Authors:** Hayley Macleod, Luisa Weiss, Sarah Kelliher, Barry Kevane, Fionnuala Ní Áinle, Patricia B. Maguire

## Abstract

Extracorporeal Photopheresis (ECP) is a leukapheresis based treatment for Cutaneous T-Cell Lymphoma, which takes advantage of the cellular lethal effects of UVA light in combination with a photoactivated drug, 8-methoxypsoralen. 25% of patients treated with ECP do not respond to treatment, however the underlying mechanisms for this lack of response remain unknown. Platelets, a rich source of extracellular vesicles (EVs) and key mediators in thromboinflammatory oncological progression, as well as leukocytes, are both processed through ECP and are subsequently transfused back into the patient, delivering potent immunomodulation. The effect of exposing platelets and their EVs directly to UVA/8-methoxypsoralen is currently unknown.

Platelet-rich plasma (PRP) was isolated from healthy donors and exposed to UVA light and/or 8-methoxysporalen *in vitro* and platelet activation and aggregation was assessed. EV size and concentration were also characterised by Nanoparticle Tracking Analysis and Flow Cytometry. We found that UVA light and 8-methoxypsoralen treatment *in vitro* did not induce platelet aggregation or alter significantly levels of the platelet activation markers soluble P-selectin or platelet factor 4, with circulating levels of small and large EV size and concentration remaining constant. Therefore, utilising the combination of UVA light and 8-methoxypsoralen used in ECP *in vitro* does not activate platelets or alter important circulating EVs. Further studies will be needed to validate if our observations are consistent *in vivo*.

## 1. Introduction

Extracorporeal Photopheresis (ECP) is a leukapheresis-based treatment developed by Richard Edelson in the 1980s to treat Cutaneous T-Cell Lymphoma (CTCL)(1). This therapeutic approach takes advantage of the cellular lethal effects of ultraviolet A light in combination with a photoactivated drug 8-methoxypsoralen on leukocytes, resulting in apoptosis of these treated cells. Reinfusion of the treated blood fraction to the patient delivers potent, personalized, immunomodulation (1). In 1988, the US Food and Drug Administration (FDA) granted approval of ECP as a palliative treatment for skin manifestations seen in CTCL, becoming the first ever FDA approved immunotherapy-based cancer treatment(2),(3). Subsequently, ECP has passed approval for clinical use in the treatment of Graft Vs Host disease, scleroderma, and solid transplant rejection(4). ECP involves a double-needle blood draw into a closed circuit. Blood centrifugation separates a specialized “buffy coat” fraction (of which contains both leukocytes and platelets)(5) from the erythrocytes and plasma. The latter are then returned to the patient in real time. The buffy coat fraction is treated with a defined dose of 8-methoxypsoralen and UVA light before transfusion back into the patient.

The exact mechanism of action of ECP is not comprehensively understood however it is proposed that, upon UVA exposure, 8-methoxypsoralen becomes activated, binding and crosslinking DNA within lymphocytes and causing apoptosis(6). Auto-antigens released during this process (initially and also post-ECP) permit priming of monocytes into specially-differentiated auto-dendritic cells (DC). These specialized DC target the autologous malignant lymphocytes for destruction post-transfusion, facilitating immunomodulation(7). The half-life of 8-methoxypsoralen is only microseconds, enabling the treated lymphocytes to retain their antigenicity to prime specialized monocytes against autoantigens with minimal adverse side effects(8, 9). This led to the first ECP clinical trial in 1987, including 37 chemotherapy resistant CTCL patients. While most patients responded to ECP, a subset of patients (27%) were non-responders, although the underlying mechanisms for non-response were unknown(2). In the years that followed, the American Council on ECP was formed to develop a consensus report on the scientific and clinical progress of ECP. It was agreed that ECP is a bidirectional therapy with a simultaneous immunizing and tolerizing effect, characterized as a dendritic antigen-presenting cell-based therapy(10). A meta-analysis review analyzing 19 studies with varying degrees of CTCL concluded the overall response rate of monotherapy ECP treatment was 55.5% with 15% of patients achieving a complete response(11). The downstream mechanisms for these clinical responses and the mechanisms underlying different response rates to ECP remain poorly characterized. It has been proposed that platelets may play a role in such mechanisms as they are key mediators in several processes highly relevant in the ECP physiological progression of inflammation, immune cell interactions, angiogenesis, tumour growth and migration(12). Many of these processes are highly relevant in ECP.

Pioneering work by Durazzo *et al* interrogated the potential contribution of platelets to ECP responses by imitating the ECP process with a chamber device, indicating that if platelets adhere to the irradiation plate, they potentially engage with monocytes in a P-selectin-dependent interaction. Intriguingly, platelet adhesion to the plate positively correlated with dendritic cells differentiation(5, 13), highlighting the potential involvement of platelets in ECP and the need for further expansion of this hypothesis. However, it is still unknown if and at what stage platelets activate throughout the ECP process. Furthermore, the effect of the specific ECP dose of UVA light and 8-methxoypsokiralen on platelets and their derived extracellular vesicles is currently unknown.

Platelets are key mediators of the immune response, expressing and releasing potent protein regulators upon activation, including macrophage inflammatory protein-1α (MIP-1α), monocyte chemotactic protein-3 and RANTES, all of which are known activators of leukocytes(14, 15). Platelets acquire and express a sub-set of T-cell co-stimulatory molecules, such as CD40, CD80, CD86, and MHC receptors, from their parent megakaryocytes, facilitating antigen-presentation, T-cell and B-cell activation(16),(17),(18). Moreover, small platelet derived extracellular vesicles (pEVs) are differentially packaged into α-granules and specifically released upon activation, with larger pEVs blebbing off the platelet plasma membrane. pEVs possess proinflammatory and immune regulatory potential through transfer of their cargo proteins including interleukin-1β(19), C-reactive protein and damage-associated molecular patterns (e.g. HMGB1)(20) to downstream target cells(21). Furthermore, pEVs can directly associate with granulocytes and monocytes within circulation(22), and express antigen loaded MHC-I on their membrane, hence pEVs are capable of initiating a specific CD8^+^ T-cell proliferation response(23). This enables platelets and their EVs to function as antigen-presenting cells *in vivo*. Collectively, these data suggest that the specific effects, functions, and activation of platelets in ECP warrants more detailed characterization in order to either consider *or, of equal importance, to rule out* this pivotal potential mechanism of action, in order to bring the field closer to understanding the *true mechanism* underlying the clinical effects of ECP. This study therefore aimed to investigate the effects of UVA light and 8-methoxypsoralen exposure used in the ECP process on platelet and circulating extracellular vesicles, to characterize their involvement in the mechanisms of action underlying the clinical effects of ECP.

## 2. Materials and Methods

### 2.1 Donor blood collection

Healthy volunteers (n=3) were recruited between 04/10/2022 and 12/10/2022 following informed consent according to the declaration of Helsinki at the Conway Institute, University College Dublin, Ireland. Blood was collected via venipuncture with a 21-gauge needle into four 10 ml Acid Citrate Dextrose-A (ACD-A) vacutainers (BD Vacutainer, Franklin Lakes, New Jersey, USA). Whole blood was centrifuged at 200 xg for 10 min (no break) to obtain Platelet Rich Plasma (PRP) for downstream experiments. For Platelet Poor Plasma (PPP), PRP (post-treatment) was further centrifuged at 2000 xg for 10 min to obtained PPP for downstream experiments. Platelet Poor Plasma (PPP) was aliquoted and stored at −80 °C until further use.

### 2.2 ECP dose UVA light and/or 8-methoxysporalen Exposure

PRP was exposed to UVA light and/or 8-methoxysporalen as per the protocol detailed in Figure1. PRP was exposed to 1.5 Joules/cm^2^ UVA light in the presence or absence of 336 ng/ml 8-methoxysporalen; 8-MOPS (UVADEX, Mallinckrodt Pharmaceuticals, Staines-upon-Thames, UK). UVA light and 8-methoxypsoralen dose stated above, calculated as per Mallinckrodt manufacture instructions(24, 25).

A UVAB light meter (RS-Pro, London, UK), which measures in mW/cm^2^, was used to ensure the correct wavelength and intensity of light was used (Supplementary Figure. 3). PRP post-exposure was subsequently used in the 96-well plate aggregometry assay. For all other downstream experiments, PRP post-exposure was further centrifuged at 2000 xg for 10min to obtain PPP.

### 2.3 96-well plate aggregometry assay

A dose response curve was obtained for the agonist adenosine diphosphate (ADP) for each donor (n=3) to ensure <50% aggregation was achieved (Supplementary Figure.1).

The 96-well plate aggregometry assay was carried out as previously described(26). PPP was used as the 100% aggregation control with PRP as the 0% aggregation control. 45 µl of sample (PRP or PPP, treated PRP) was plated with 5 µl of agonist (1.25 µM ADP; JNL buffer for controls), onto a half-area flat-bottom 96 well plates (Greiner Bio-One, Frickenhausen, Germany) and incubated for 5min at 1200rpm at 37°C. Agonist was added prior to or post UVA exposure to evaluate the activated compared to resting aggregometry effects (Figure 1). Results were read on a Clariostar plate reader (BMG Labtech, Ortenberg, Germany) under the spiralled light capture setting, at a wavelength of 575nm.

**Figure 1.**
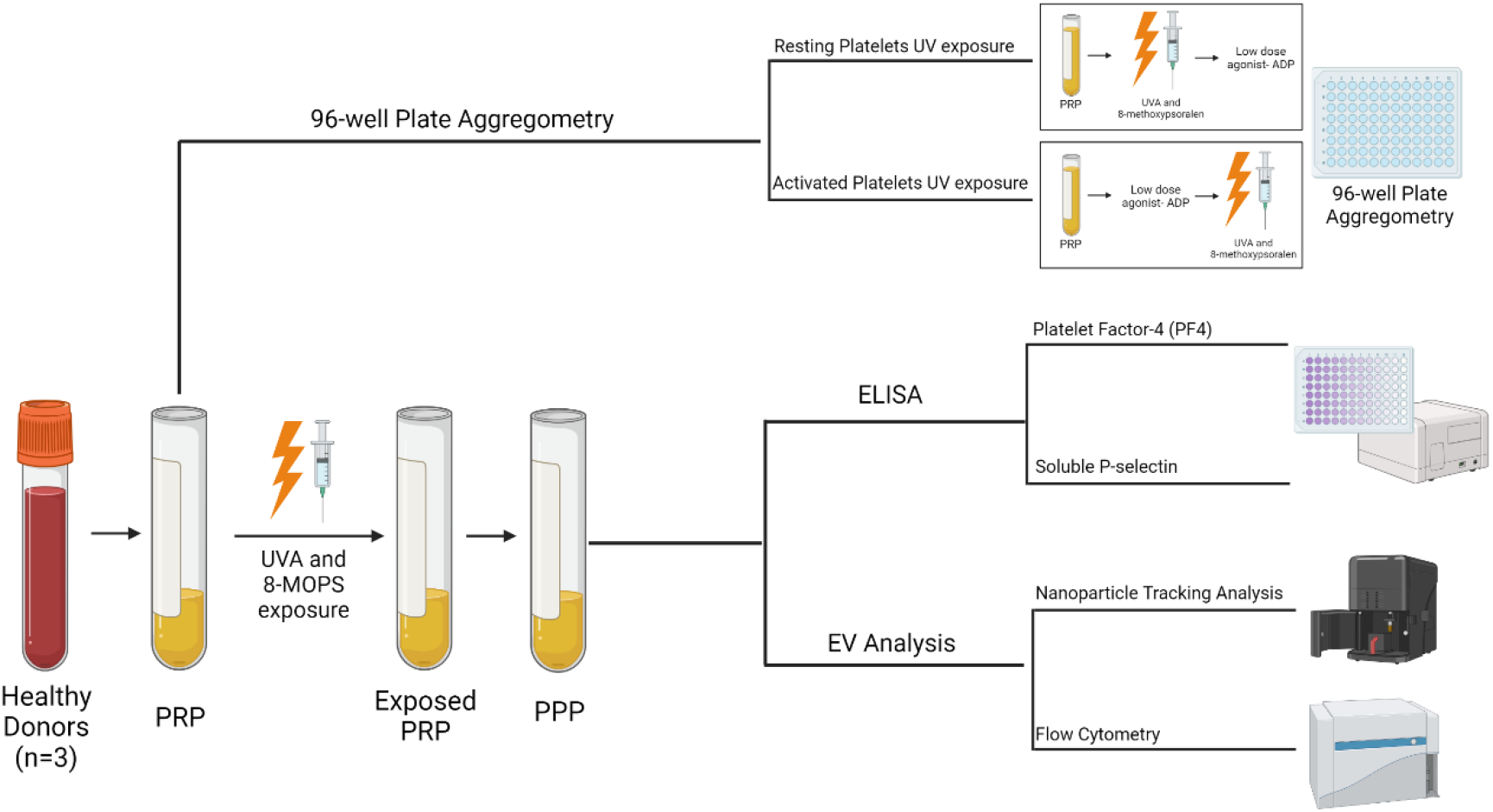
The methodology investigating the effect of UVA and 8-methoxypsoralen (8-MOPS) on platelets and EVs. Healthy volunteers (n=3) blood was collected via venipuncture into four 10 ml Acid Citrate Dextrose-A (ACD-A) vacutainers. Whole blood was centrifuged at 200xg for 10 min (no break) to obtain Platelet Rich Plasma (PRP). To access platelet aggregation, a 96-well plate aggregometry assay was performed under resting and activated platelet conditions. PRP was exposed to 1.5 Joules/cm^2^ UVA light in the presence or absence of 336 ng/ml 8-methoxysporalen, dose stated above calculated as per Mallinckrodt manufacture instructions(24),^(25)^. For resting conditions, PRP was exposed to UVA/8-methoxypsoralen before activating with 1.25 µM ADP, where-as for activated conditions, PRP was activated with 1.25 µM ADP followed by exposure to UVA/8-methoxypsoralen. 45 µl of sample (PRP or PPP, treated PRP) was plated with 5 µl of agonist (1.25 µM ADP; JNL buffer for controls), onto a half-area flat-bottom 96 well plates and incubated for 5min at 1200rpm and 37°C. PPP was used as the 100% aggregation control with PRP as the 0% aggregation control. Results were read on a Clariostar plate reader under the spiralled light capture setting, at a wavelength of 575nm. PRP post-exposure was further centrifuged at 2000xg for 10min to obtained PPP for downstream ELISA and EV analysis. Sandwich ELISAs for P-selectin and platelet factor 4 were performed according to the manufacturer’s instructions. Small EV size and concentration was measured with Nanoparticle Tracking Analysis using a NanoSight NS300 system. PPP was diluted (1:100–1:500) in PBS to achieve 10 to 60 particles per frame, according to the manufacturer’s instructions. Sample analysis was performed at 25°C and a constant flow rate of 50. The 15 × 60 s videos were captured with a camera level of 13 and data were analysed using NTA 3.1.54 software with a detection threshold of 10. Flow cytometry analysis of circulating large EVs was performed using a CytoFlex S with particle size calibrated using commercially available polystyrene beads. Compensation for differences in refractive indices was performed using Rosetta calibration beads (Exometry) and the Rosetta calibration software (v1_30). 100-, 300-, 500-, and 900-nm analysis gates were established. Data analysis was performed using Kaluza. The 30 µL platelet-poor plasma (PPP) was diluted with 520 µL 0.22 µm filtered PBS. Samples were further diluted 1:20 to prevent EV swarming and analysed in triplicate at a constant flow rate of 10 µL/min for 2 min or until 100,000 events were recorded. A buffer-only (0.22-µm filtered PBS) sample was assayed using the same settings and during the same experiment as the samples and background vesicle counts were subtracted from the respective samples. 8MOPS: 8-methoxypsoralen.

### 2.4 ELISA

ELISAs were used to determine platelet activation markers in corresponding PPP. Sandwich ELISAs for P-selectin (CD62P, R&D Systems, Minneapolis, Minnesota, USA) and platelet factor 4 (PF4, R&D Systems) were performed according to the manufacturer’s instructions. All standards and samples were assayed in duplicate.

### 2.5 Nanoparticle tracking analysis

Particle size distribution in PPP was determined by NTA using a NanoSight NS300 system (Malvern Technologies, Malvern, UK) fitted with a 488-nm laser and a high-sensitivity scientific camera, as previously described(27). Plasma was diluted (1:100–1:500) in particle-free phosphate-buffered saline (PBS; Gibco, Waltham, Massachusetts, USA) to achieve 10 to 60 particles per frame, according to the manufacturer’s instructions. Sample analysis was performed at 25°C and a constant flow rate of 50(27). The 15 × 60 s videos were captured with a camera level of 13 and data were analysed using NTA 3.1.54 software with a detection threshold of 10 as documented before(27).

### 2.6 Flow cytometry

Flow cytometry analysis of circulating EVs was performed using a CytoFlex S (Beckman Coulter, Brea, USA), with particle size calibrated using commercially available polystyrene beads. Polystyrene beads are solid spheres with a refractive index of 1.61(28), whereas vesicles consist of a core and a shell with refractive indices of 1.38 and 1.48, respectively. Compensation for differences in refractive indices was performed using Rosetta calibration beads (Exometry, Amsterdam, Netherlands) and the Rosetta calibration software (version 1_30) (28). This calibration is based on the Mie theory and used to relate side scatter intensities (in arbitrary units) to EV diameter (in nm) for a given refractive index(29). 100-, 300-, 500-, and 900-nm analysis gates were established. Data analysis was performed using Kaluza (version 2.1, Beckman Coulter). The 30 µL platelet-poor plasma (PPP) was diluted with 520 µL 0.22 µm filtered PBS. Samples were further diluted 1:20 to prevent EV swarming and analysed in triplicate at a constant flow rate of 10 µL/min for 2 min or until 100,000 events were recorded. A buffer-only (0.22-µm filtered PBS) sample was assayed using the same settings and during the same experiment as the samples and background vesicle counts were subtracted from the respective samples.

### 2.7 Statistical analysis

Statistical analysis of differences in aggregation, protein expression and EV levels were assessed in RStudio. Data were tested for normal distribution using a Shapiro-Wilk Test. Normally distributed data were assessed for statistical significance using a one-tailed ANOVA test. Non-normally distributed data was assessed for statistical significance using a Kruskal-Wallis test. P-values < 0.05 were regarded statistically significant.

## 3. Results

### 3.1 ECP dosage UVA light/8-methoxypsoralen treatment does not activate platelets

Upon activation, platelets release soluble proteins that subsequently elicit downstream effects, including angiogenesis, inflammation and immune regulation (12). Platelet Factor 4 (PF4) is a potent activation marker contained within α-granules that is rapidly released upon platelet stimulation. Furthermore, P-selectin is expressed on the plasma membrane upon activation, and is subsequently cleaved, releasing its soluble form into circulation(30). Circulating levels of these protein markers correlate with platelet activation in various conditions(31). To explore the potential dual effect of UVA light and 8-methoxypsoralen exposure on platelet activation, immunoassays for both PF4 and soluble P-selectin were performed in corresponding PPP. No significant difference was observed in α-granule-component release, quantified by PF4 levels between groups (*p*= 0.141, Figure. 2A). Similarly, soluble P-selectin levels quantified remained unchanged (*p*= 0.918, Figure. 2B), suggesting that the UVA light and/or 8-methoxypsoralen at doses used during ECP do not activate platelets *in vitro*.

**Figure 2.**
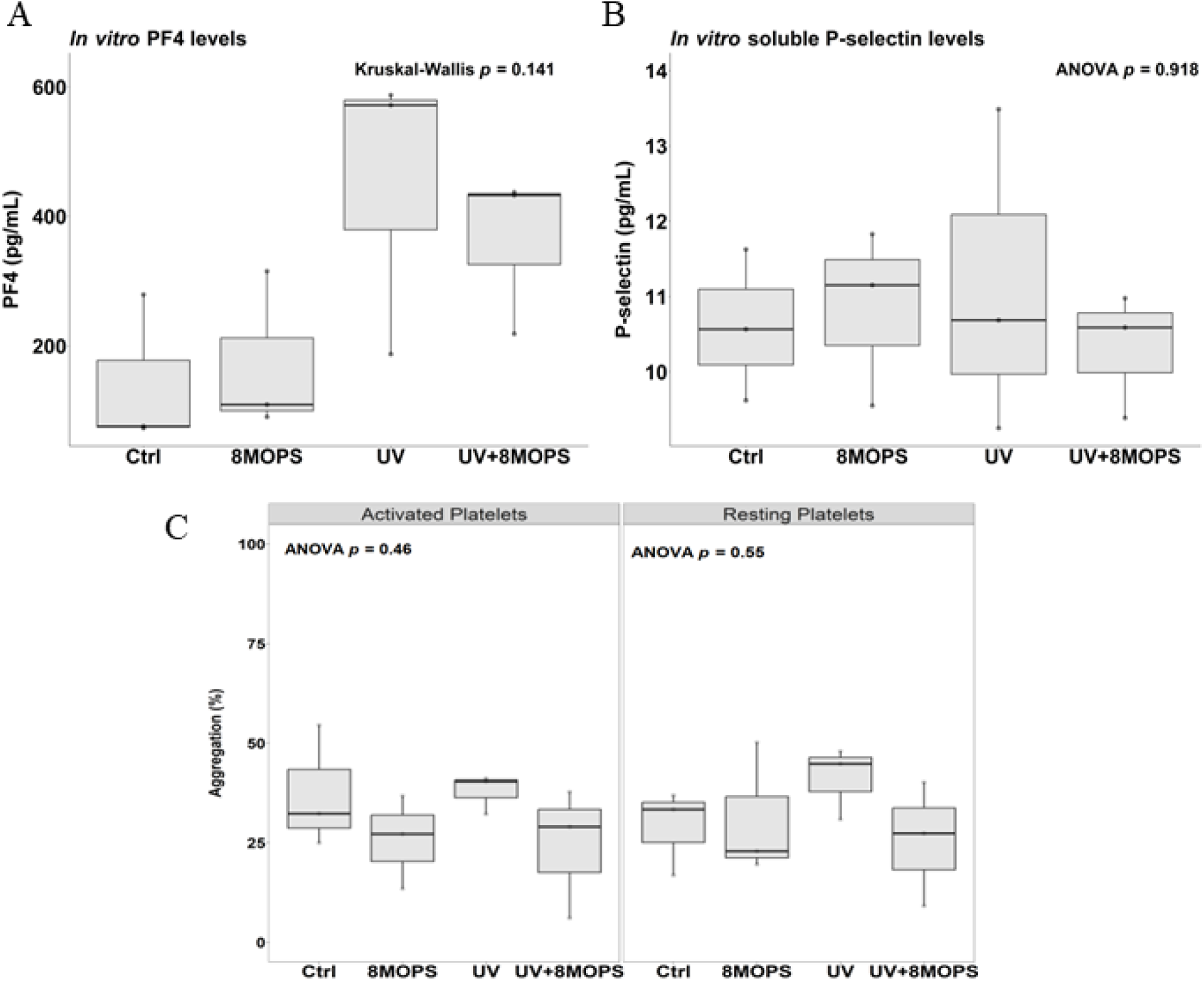
ECP dose UVA light/8-methoxypsoralen (8-MOPS) treatment does not activate platelets. Platelet activation markers, PF4 and soluble P-selectin were quantified using immunoassays in PPP prepared from PRP following exposure to UVA light and/or 8-methoxypsoralen. (A) PF4 (Kruskal-Wallis test, *p*=0.141) and (B) soluble P-selectin levels (one-way ANOVA, *p*=0.918) did not differ following UVA/8-methoxypsoralen exposure. (C) Platelet aggregation was performed using a 96-well plate aggregation assay, with PRP exposed to UVA light and/or 8-methoxypsoralen in a resting state as well as ADP-activated state (mid-range stimulatory dose of ADP to achieve first wave of platelet activation (<50% aggregation). UVA/8-methoxypsoralen treatment did not affect platelet aggregation across activation states (activated platelets; *p*=0.46; resting platelets; *p*= 0.55). 8MOPS: 8-methoxypsoralen.

Next, we assessed whether UVA light/ 8-methoxypsoralen exposure primes platelets for activation post-ECP treatment. Notably, platelet activation occurs in two waves, the first as a direct result of a stimulus, facilitating platelet adhesion, granule secretion and aggregation, followed by a second wave, amplifying platelet activation and aggregation through positive feedback regulation(30). PRP was activated using a donor-specific mid-range stimulatory dose of ADP to achieve this first wave of platelet activation (<50% aggregation, Supplementary Figure 1). Mild platelet stimulation enabled the aggregation capacity of UVA/8-methoxypsoralen treatment to be investigated. ADP-activated and resting platelets were exposed to a UVA dosage of 1.5 J/cm^2^ and/or 336 ng/ml of 8-methoxypsoralen. As seen in Figure 2C, UVA with or without 8-methoxypsoralen treatment did not induce platelet aggregation under either activated or resting conditions (*p*=0.46 and *p*=0.55, respectively), highlighting that 1.5 J/cm^2^ UVA light and 8-methoxypsoralen treatment do not prime, stimulate, or increase platelet activation *in vitro*.

### 3.2 Extracellular vesicles (EVs) release remains constant upon ECP dose UVA/8-methoxypsoralen exposure

Nanoparticle tracking analysis was used to determine the concentration and size of small EVs (50-200nm) between treatment conditions. Total particle counts (*p*= 0.988, Figure. 3A) and particle mode size (*p=* 0.439, Figure. 3B) remained unchanged, between the UVA/8-methoxypsoralen treatment and individual control conditions. Flow cytometry analysis of large EVs between 100–1000 nm also revealed no difference in either total particle count (*p* = 0.964, Figure 3C) or vesicle size between treatment groups (Supplementary Figure. 2).

**Figure 3.**
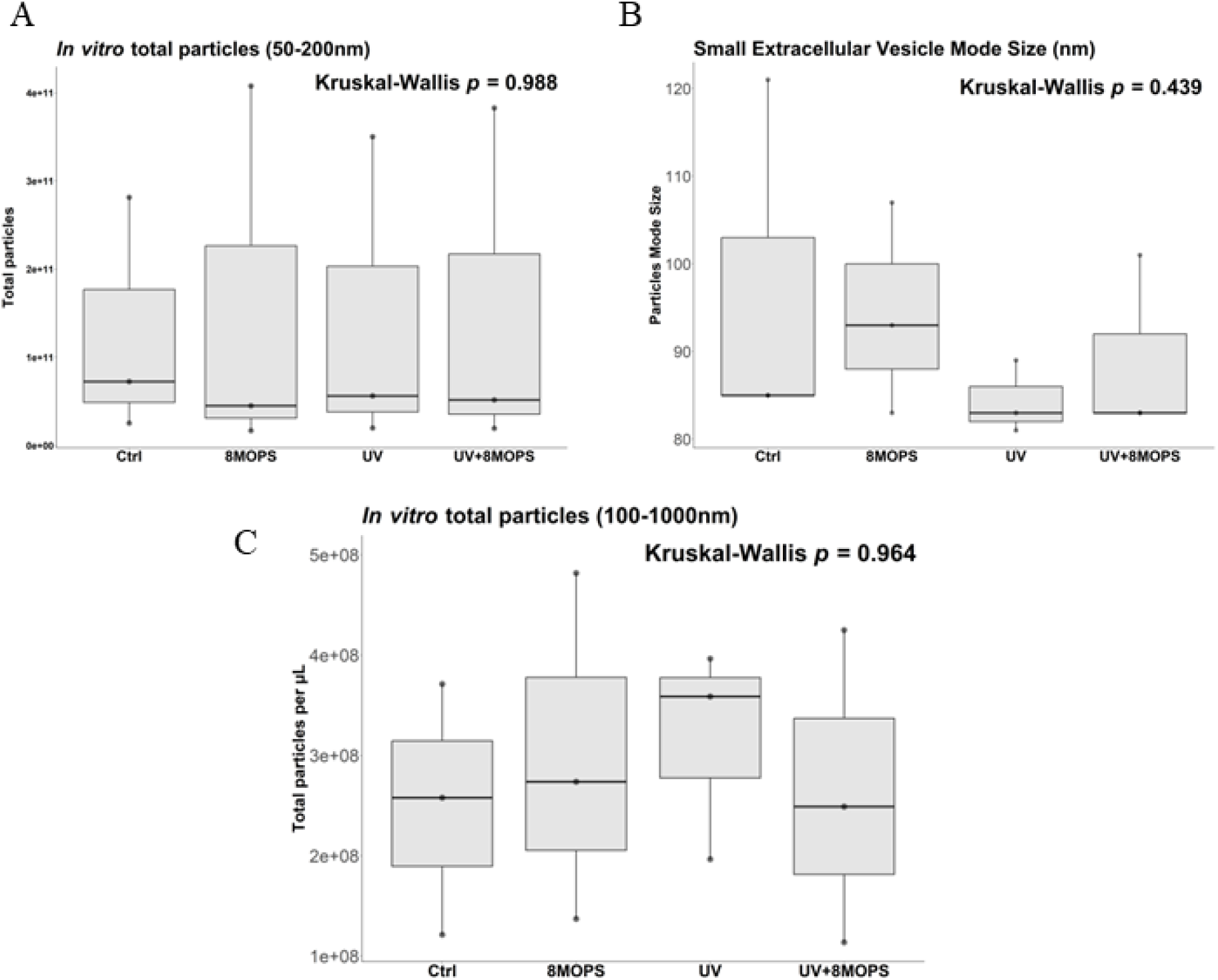
Extracellular vesicle release upon ECP dose UVA /8-methoxypsoralen treatment remains unchanged. Vesicle quantification and sizing of circulating small EVs (50-200nm) in PPP prepared from PRP following exposure to UVA light and/or 8-methoxypsoralen was assessed using Nanoparticle Tracking Analysis (NTA) with the Nanosight NS300. 15 × 60 s videos were captured at a camera level of 13 and analysed using a detection threshold of 10. (A) Small EV concentration (Kruskal-Wallis test, *p=* 0.988) and (B) size (Kruskal-Wallis test*, p=* 0.439) remained unchanged post-treatment. (C) Flow Cytometry was performed with the CytoFlex S to quantify large EVs (100-1000nm) in PPP post-exposure to UVA light and/or 8-methoxypsoralen. Light scatter intensities were adjusted to reflect biological EV properties using Rosetta calibration beads and software. Large EV concentration (Kruskal-Wallis test, *p=* 0.964) remained unchanged upon dual and individual UVA light/8-methoxypsoralen exposure. 8MOPS: 8-methoxypsoralen.

Collectively, characterization of the small and large EVs highlight that UVA light/ 8-methoxypsoralen exposure does not influence extracellular vesicle release or size distribution. This low dose UVA light/8-methoxypsoralen treatment used in ECP does not alter important circulating EVs, which is important to understand, as this treated blood fraction is transfused back into the patient post-treatment.

## Discussion

We have shown that *in vitro* exposure of platelets to doses of UVA light/8-methoxypsoralen utilized during ECP does not affect platelet activation, aggregation or alter circulating extracellular vesicle count or size. Elucidating the platelet contributions to the mechanism of action underlying the clinical effects of ECP is a crucial step towards revealing the true mechanism, and towards harnessing this knowledge to understand fluctuating clinical response rates and optimizing clinical benefit.

In line with historical studies investigating the effect of UVA/8-methoxypsoralen on platelet function (32–36), we have shown that UVA light and 8-methoxypsoralen treatment does not alter platelet activation or circulating EV profiles. There are few published data characterizing the effects of ECP on platelets, however several studies that investigated platelet activation in response to similar treatments are in line with our findings. Historically, UVA light was coupled with 8-methoxypsoralen at various doses (and other psoralen derivatives) for dermatological treatments and was termed Psoralen Ultra-Violet A (PUVA) therapy. This was a common treatment for vitiligo and psoriasis, in which topical and later systemic 8-methoxypsoralen treatment was administrated to the affected patient followed by UVA exposure to the skin(37). In the early 90s, Procaccini *et al.* demonstrated no abnormality in the platelet aggregation patterns with ADP stimulation (or other agents including collagen, ristocetin and arachidonic acid) using conventional aggregometry assays on PRP treated with 200 ng/ml of 8-methoxypsoralen and/or 5 J/cm^2^ UVA exposure in healthy individuals. Furthermore, in an *ex vivo* study, the authors showed the platelet aggregation profiles were not altered after 8-methoxypsoralen ingestion or PUVA treatment. Platelet aggregometry was performed on PRP at basal conditions, 2.5 hours after oral ingestion of 8-methoxypsoralen (0.6-0.8 mg/kg) and after 4 PUVA sessions. None of these 4 patients showed modification to their platelet aggregation profile after either 8-methoxypsoralen ingestion or PUVA treatment(38). Similarly, Rao *et al.,* showed patients exposed to PUVA therapy (20mg of 8-methoxypsoralen and 7.8 mW/cm^2^ for a total of 10 joules) for the treatment of vitiligo, did not show any alterations in arachidonic acid-induced platelet activation or prostaglandin synthesis(39). In the efforts to elucidate the mechanisms underlying PUVA therapy the phospholipid mediator platelet-activating factor (PAF) pathway was knocked out in a mouse model, with subsequent treatment with 4.5 mW/cm^2^ UVA light and 1 mg/ml 8-methoxypsoralen. The PAF pathway was found to be crucial for PUVA-induced immune suppression, playing a role in skin inflammation and apoptosis(40). Therefore, there is a potential that platelets are not directly activated from 8-methoxypsoralen/ UVA exposure but through the PAF pathway post treatment. However, this hypothesis needs to be investigated in the context of ECP.

Furthermore, UVA irradiation of platelets has been used for bacterial sterilisation for over 30 years. Lin *et al*, examined the effects of long-wavelength UV energy (3.5 to 4.8 mW/cm^2^ UVA light) in combination to 8-methoxypsoralen (300 µg/mL) on platelet concentrates for bacterial and viral inactivation^39^. In line with our results, platelet function and quality remained intact post treatment, with no significant differences in morphology scores, platelet yields or LDH levels post radiation/drug exposure. Calcium ionophore agonist A23187-induced platelet aggregation was performed 18 hours after irradiation, giving a comparable response between treated and untreated control platelets. There was also no significant difference in the thromboxane B-2 release or alpha/dense granule secretion between conditions(32), mirroring our observations that circulating α-granule markers and pEV levels remained unaltered. Subsequent investigations echoed these results, indicating the safe use of UVA light and 8-methoxypsoralen to sterilise platelet concentrates for blood transfusions(33, 41). Of interest to this current study, Grass *et al.* demonstrated leukocytes are inactivated in platelet concentrates used for blood transfusion, through treatment with UVA light and 8-methoxypsoralen at incrementally increasing doses. Results indicate the utility of such a procedure has the potential to reduce the incidence of leukocyte-mediated adverse immune reactions, while retaining platelets quality for transfusion(42). Moreover, the INTERCEPT clinical platelet sterilisation method (3 J/cm^2^ UVA and 150 μmol/l amotosalen treatment) utilises ultraviolet radiation for pathogen inactivation of platelet concentrates(34). This UV treatment does not influence platelet *in vitro* function and quality(35), retaining *in vivo* platelet survival and consistent platelet transfusion recovery rates(36). Collectively, these results highlight the low dose UVA light and 8-methoxpsoralen used in ECP does not activate, aggregate or influence platelets upon exposure.

In contrast, the results from our study contradict a recent publication by Budde *et al*. Here, authors investigated the effect of UVA light and 8-methoxypsoralen on red blood cells, platelets and the generation of reactive oxygen species, revealing platelets to be highly activated after 8-methoxypsoralen and UVA treatment(43). The divergence with our results may have arisen from differences in the fundamental underlying methodology. Specifically, the authors exposed a buffy coat fraction containing a mixture of white blood cells and platelets to 2 J/cm^2^ UVA light and 200 ng/mL 8-methoxypsoralen, therefore the activation results observed may not be due to the effect of this exposure to platelets alone but instead from a combination of interactions between the activated leukocytes and platelets(43). It has been shown that ECP treatment causes apoptosis of exposed lymphocytes, however 40% of treated cells retain their cellular integrity(44),(41). Furthermore, monocytes are resistant to ECP-inducted apoptosis, facilitating phagocytosis of apoptotic lymphocytes post-exposure, maturing into antigen-presenting dendritic cells(5, 41, 45). This process activates antigen presenting cells, presenting self-antigens to target malignant T-lymphocytes. These active and intact leukocytes could interact with platelets causing downstream activation(46, 47).

The result from Badde *et al* may however be essential in context of our study, as we can hypothesize that although the UVA/8-methoxypsoralen treatment does not have a direct effect on platelets, it may alternatively cause indirect activation through interactions with the exposed and subsequently activated leukocytes. This hypothesis could reveal a potential link in the mechanism of immunomodulation occurring post-ECP treatment, with further investigations into the platelet-leukocytes interactions urgently warranted to elucidate the mechanism underlying ECP.

Further *in vivo* studies are required to elucidate the exact nature of platelet activation during ECP. Platelets are very sensitive to environmental stimuli such as temperature and shear. Thus, it may be possible that platelets are activated at alternative points within the ECP process, facilitating downstream interaction with leukocytes and release of potent soluble mediators. Furthermore, extracellular vesicles are released from platelets upon activation; however, they are also released from a multitude of cells including lymphocytes, dendritic cells, monocytes and tumour cells(48). This study is the first to investigate extracellular vesicles in the context of ECP, which could reveal potent insights into immune signaling and mechanisms at play. EVs carry potent stimulatory and regulatory cargo to downstream targets, for multidirectional cellular communication. As previously discussed, dendritic cells have an integral role in ECP immunomodulation through presentation of self-antigens to initiate an immune response targeting malignant cells. Moreover, crosstalk between tumors and dendritic cells is mediated by extracellular vesicles secreted from both cell types, with dendritic cell EVs capable of stimulating other immune cells and possess the ability to promote tumor antigen-specific responses. Therefore, it is essential to uncover the role of EVs in ECP related immune mechanism and interactions.

In this *in vitro* study we have shown that UVA light and 8-methoxypsoralen alone or in combination do not activate or aggregate platelets. EV release and size were also found to remain unchanged under such exposure conditions, as they are only released from platelets upon activation. However, the ECP process is highly complex with intricate personalised mechanism at play upon individual exposure. Therefore, further *ex vivo* studies are essential to understand how the platelets respond to mechanical forces exerted within ECP, the effects of platelet-leukocyte interactions on platelet activation as well as the extracellular vesicle profiles post ECP treatment.

## Acknowledgements

We acknowledge the excellent assistance of colleagues in Mallinckrodt Pharmaceuticals. This study was supported by an investigator-led basic research award from Mallinckrodt Pharmaceuticals to PM and FNA. All participants gave informed written consent according to the declaration of Helsinki.

## Supplementary Figures

**Supplementary Figure 1.**
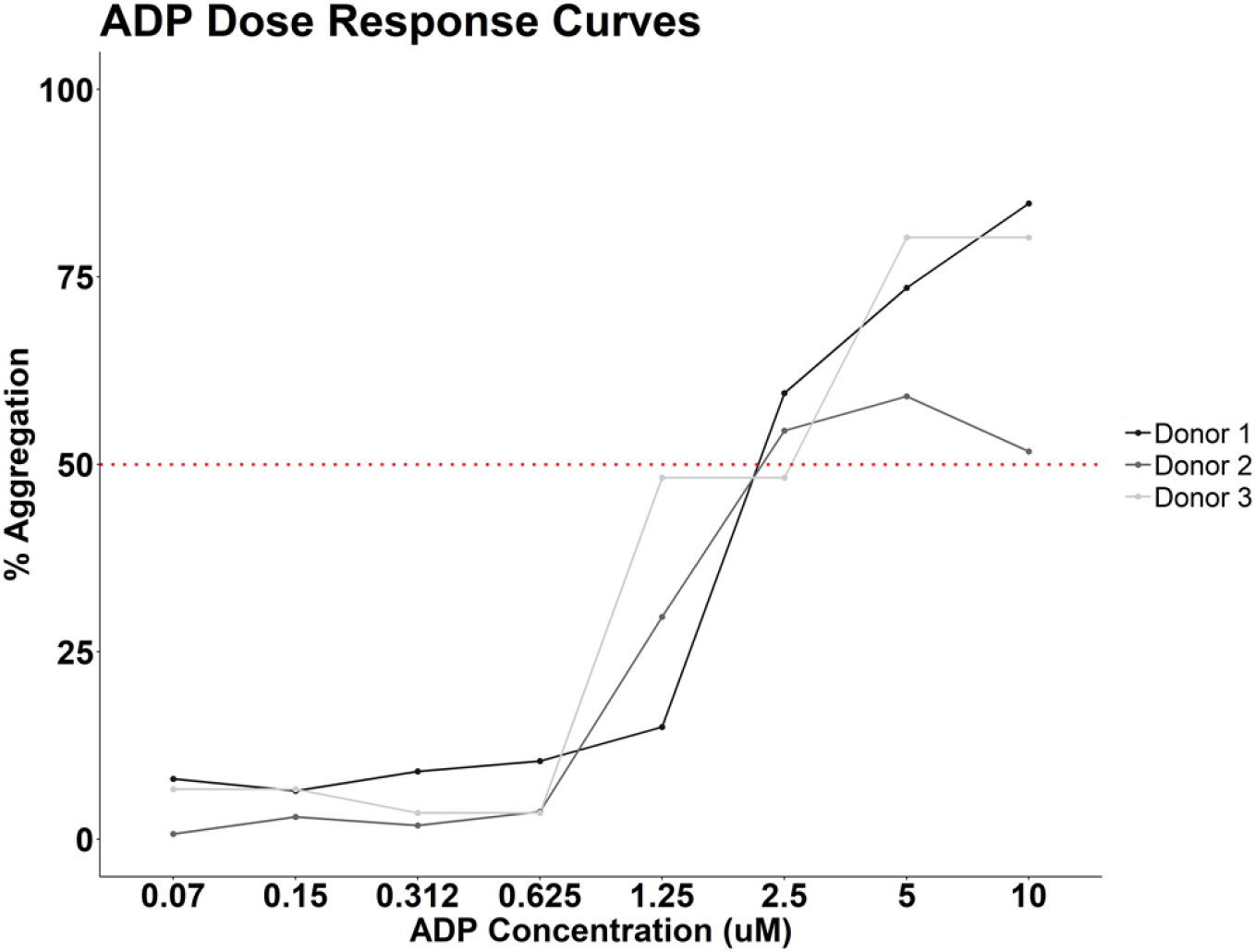
ADP Dose Response Curves. PRP was isolated from 3 healthy volunteers to carry out an adenosine diphosphate (ADP) dose response curve using the 96-well plate aggregometry assay. ADP concentrations from 0.07 µM to 10 µM ADP were assessed. 1.25 µM ADP resulted in platelet aggregation <50% aggregation (marked with a red dotted line) and was used for all follow-on experiments.

**Supplementary Figure 2.**
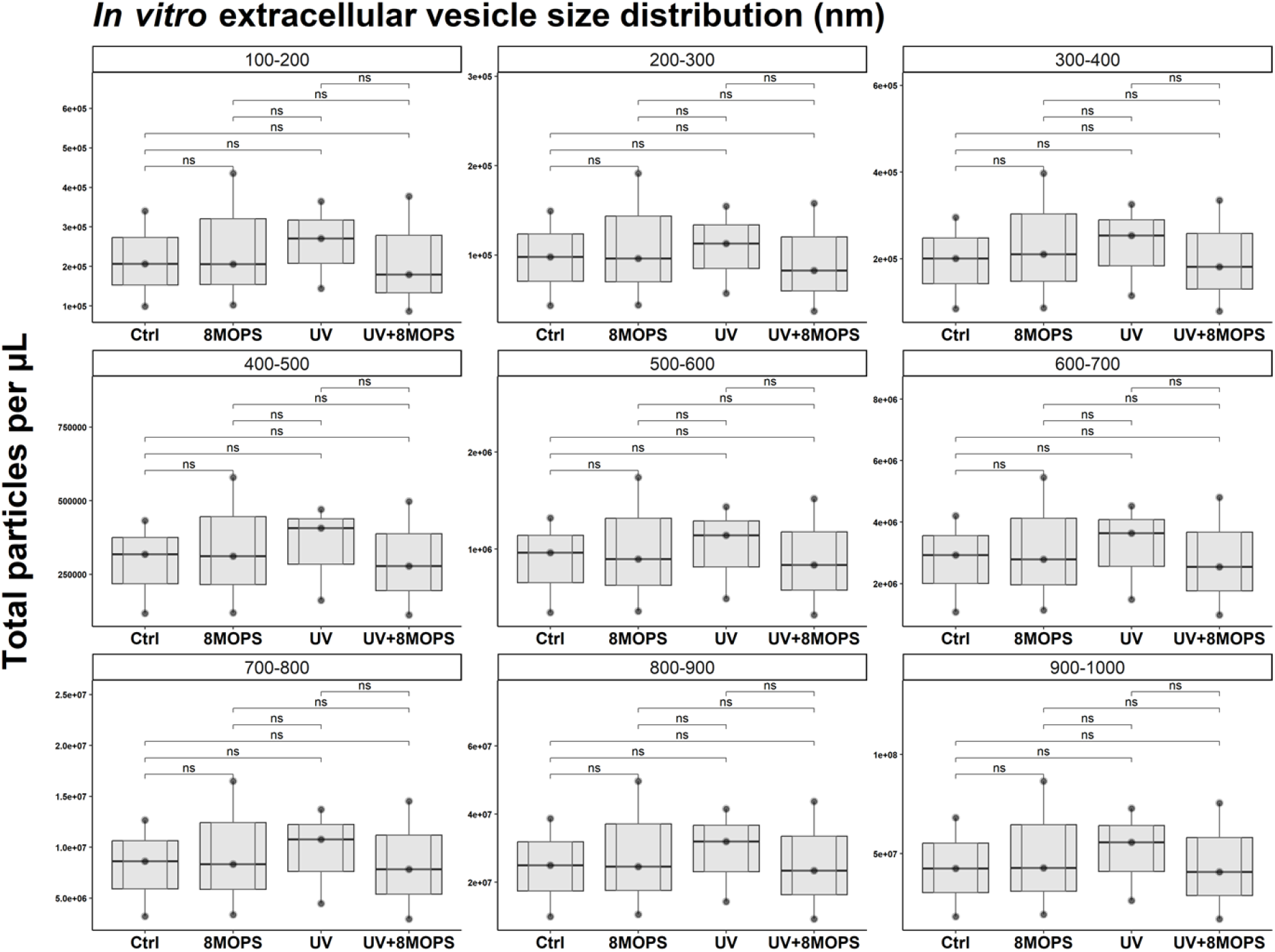
Large extracellular vesicle size remains unchanged upon ECP dose UVA /8-methoxypsoralen treatment. Flow Cytometry was performed with the CytoFlex S to quantify large EVs (100-1000nm) in PPP post-exposure to UVA light and/or 8-methoxypsoralen. Light scatter intensities were adjusted to reflect biological EV properties using Rosetta calibration beads and software. Large EV size remained unchanged upon dual and individual UVA light/8-methoxypsoralen exposure. 8MOPS: 8-methoxypsoralen.

**Supplementary Figure 3.**
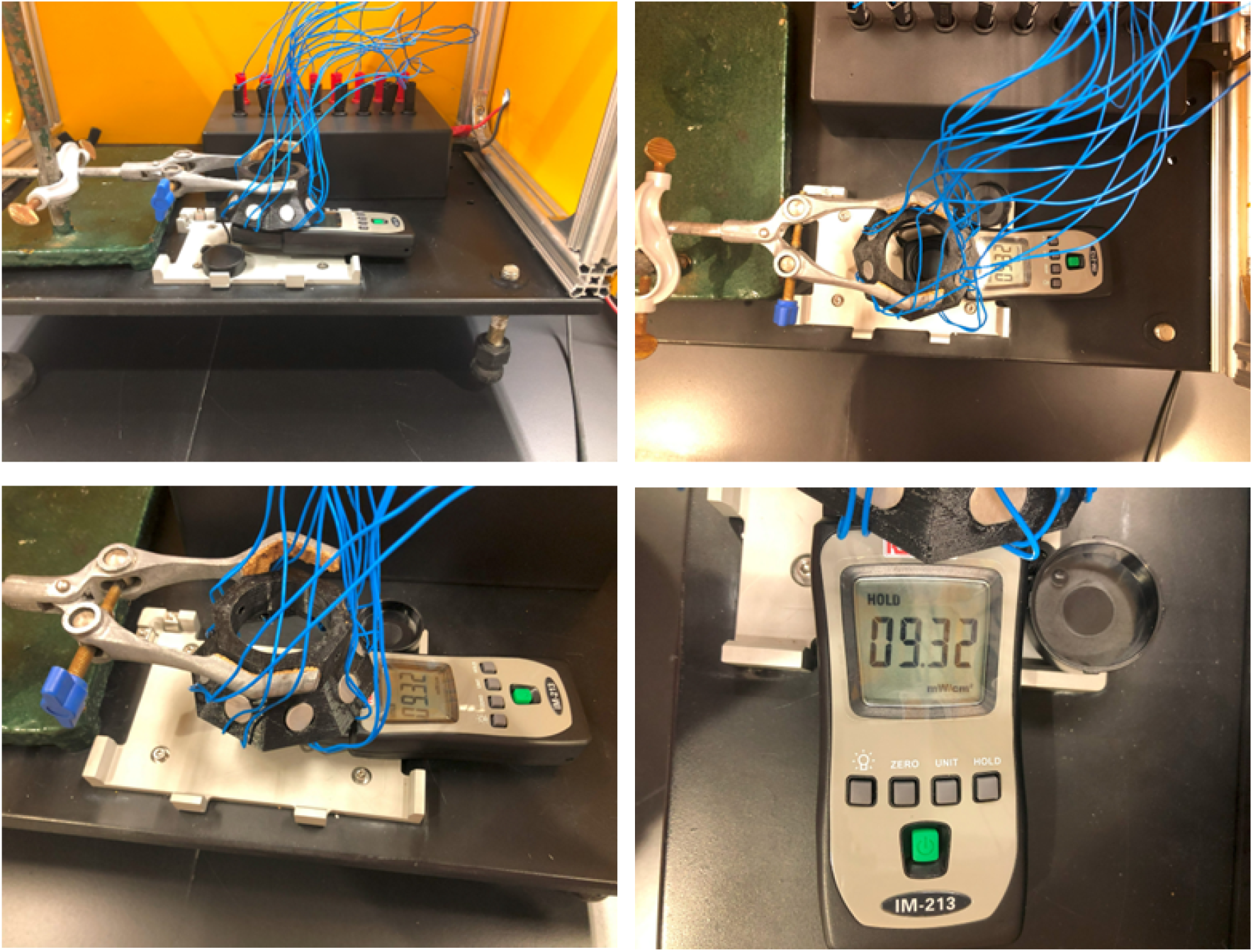
UVA light wavelength and intensity used matches the ECP UVA dose. A UVAB light meter (mW/cm^2^) was used to ensure the correct wavelength and intensity of light used to echo the ECP dose. The RS Pro UVAB light meter measures UV light in the range of 290-390nm, encapsulating the ECP UVA light range of 320-400nm. An average of 10mW/cm^2^ UV intensity was recorded at the lowest stage hight of this UV light box and using the equation Watt x Time = Joules gives you 2 min and 30 sec of this UV intensity to supply 1.5 Joules/cm^2^ UVA light to the samples in a 96-well plate.

